# Identification of candidate therapeutics and signaling pathways for multiple myeloma

**DOI:** 10.1101/2021.11.03.467084

**Authors:** Hanming Gu

**Affiliations:** SHU-UTS SILC School, Shanghai University, Shanghai, China

**Author notes:** Corresponding author: Hanming Gu, SHU-UTS SILC School, Shanghai University, Shanghai, China.

## Abstract

Multiple myeloma (MM), a plasma cell malignancy, is related to critical morbidity due to end-organ destruction. A number of factors affect the MM cell proliferation and functions. Though MM is not curable, novel targets and inhibitors have shown great effects on MM patients. Here, we aim to identify significant genes and signaling pathways of MM with SI2 treatment using a bioinformatics method. The GSE156871 dataset was originally produced by using the high-throughput BGISEQ-500. The KEGG and GO results suggested that biological pathways such as “the complement and coagulation cascades” and “the transcription activator activity” are mostly affected in the SI2 treatment of MM cells. Moreover, we identified several genes including SRC, KNG1, and PI3KCG were involved in the treatment of MM cells. Therefore, our study provides further insights into the treatment of MM.

## Introduction

Multiple myeloma (MM) is a serious malignancy which is characterized by the large expansion of malignant plasma cells in bone marrow^1,2^. MM accounts for about 15% of hematological malignancies in the worldwide^3^. The incidence of multiple myeloma is higher in African Americans, with the diagnosed age above 55 years in the US^4^. With the progress of therapeutic strategies, the median survival has improved in the past two decades^5^. The development of MM is related to the disorder of bone marrow microenvironment^6^. The MM cells in bone marrow have various interactions with immune cells and receive multiple signals that regulate the apoptosis and autophagy^6^. However, some MM cells are not dependent on the bone marrow microenvironment that leads to the secondary plasma leukemia^7, 8^. Although critical therapies have been made, most MM patients relapse with drug resistance^9^.

One of the reasons underlying drug resistance is related to the genetic abnormalities and epigenetic aberrations^10^. It has been reported that the disorder of histone methyltransferases and demethylases have been found in patients with several mutations, such as multiple myeloma SET domain^11^. This domain encodes a histone methyltransferase that regulates histone dimethylation to control the chromatin structure and change the status of gene transcription^12^. Steroid receptor coactivator-3 is amplified in various cancers including MM, which promotes many aspects of cancer such as progression and signaling pathways^13^.

In this study, we evaluated the efficacy of a newly developed SRC-3 inhibitor (SI2) by analyzing the RNA sequence data. We identified several DEGs and the relevant biological process of MM underling the SI2 treatment. We also performed the functional enrichment and protein-protein interaction (PPI) for finding MM nodes during the SI2 treatment. These important genes and pathways could be critical to improve the MM therapy and inhibit the drug resistance.

## Methods

### Data resources

The dataset GSE156871 was downloaded from the GEO database (http://www.ncbi.nlm.nih.gov/geo/). The data was produced by using anBGISEQ-500 (Homo sapiens) (Tianjin Medical University, 22 Qxiangtai Rd, Tianjin, 300070, China). The GSE156871 dataset contained four samples from the LP-1 multiple myeloma cells with the inhibitor of SRC-3 (SI-2, 25 nM).

### Data acquisition and preprocessing

The raw microarray data between wild type control samples and SI2 treatment samples were subsequently performed by R package. A false discovery rate (FDR)< 0.05 and fold change ≥1.5 were used to determine the threshold of the P-value in multiple tests. Gene Ontology (GO) and pathway enrichment analysis

The DEGs were submitted to DAVID (Database for Annotation, Visualization and Integrated Discovery, http://david.abcc.ncifcrf.gov/) for functional analysis. The enrichment analysis is based on Gene Ontology (GO, http://geneontology.org/) and Kyoto Encyclopedia of Genes and Genomes (KEGG; http://www.genome.jp/kegg/kegg2.html). Gene Ontology (GO) is divided into three categories including molecular function (MF), cellular component (CC) and biological process (BP). P< 0.05 and gene counts >10 were considered statistically significant. Module analysis

The Molecular Complex Detection (MCODE) from Cytoscape software (Cytoscape, US) was used to perform the densely connected regions in PPI networks^14, 15^. The significant changed modules were from the constructed PPI network using MCODE. The pathway enrichment analyses were then performed by using Reactome (https://reactome.org/), and P< 0.05 was used as the cutoff criterion.

## Results

### Identification of DEGs between the control and SI2 treatment MM cells

To determine whether SI2 therapy is effective in MM cells, we compared the wild type of MM cells (LP-1 cells) and the MM cells treatment with SI2 (inhibitor of SRC-3). A total of 385 genes were identified to be differentially expressed in SI2 treatment group with the threshold of P<0.05.

The heatmap and volcano plot indicate that DEGs were up-regulated and down-regulated in SI2 treatment from the control group in the MM cells (Figure 1). DEGs were also showed in volcano plot, with 20 up-regulated and down-regulated genes annotation. The selected top 10 up- and down-regulated genes are listed in Table 1.

**Figure 1.**
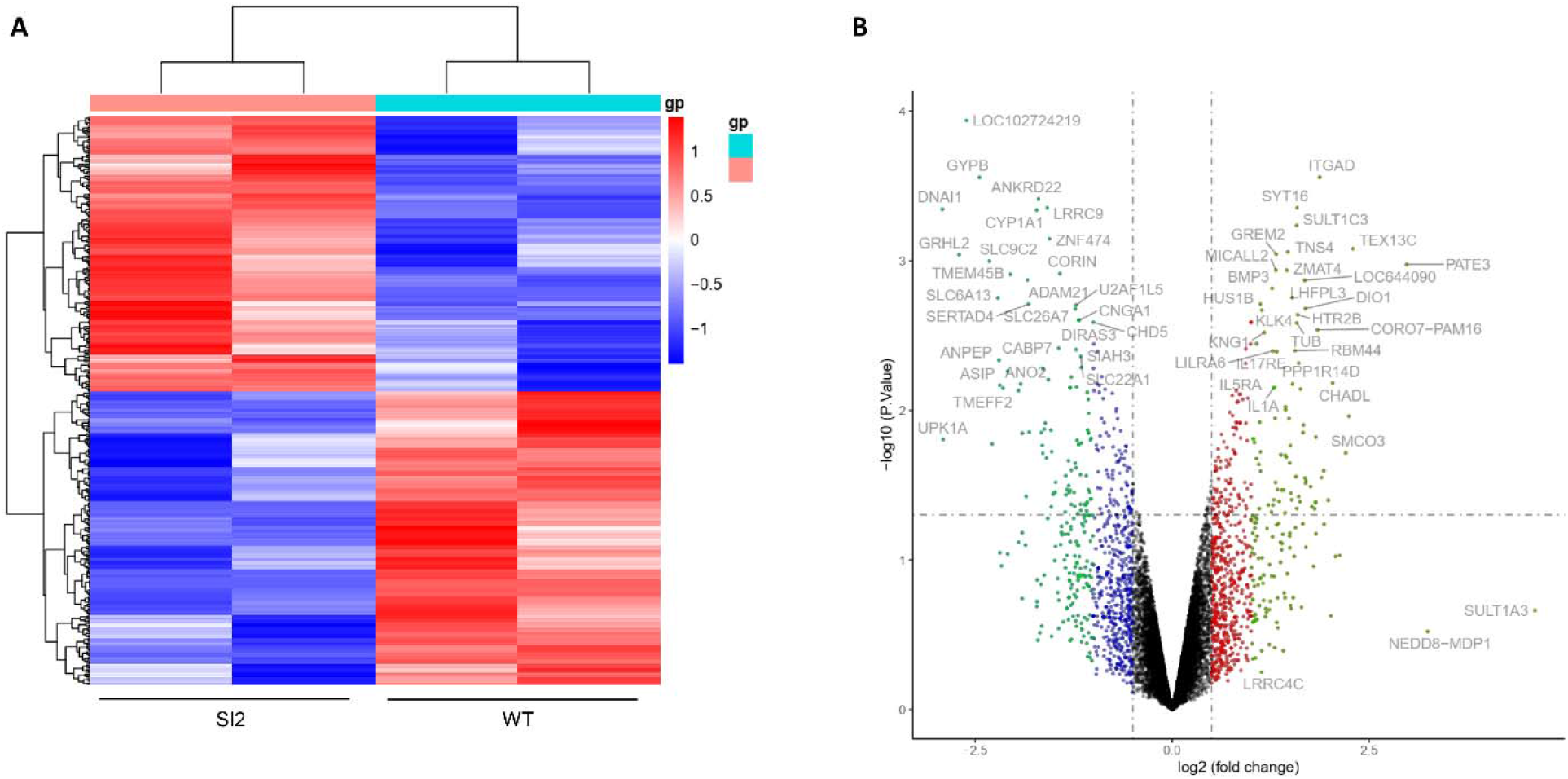
Heatmap and volcano plots of differential expression. (A) Heatmap shows 384 significantly (FDR ≤ 0.05) differentially expressed genes between WT and Si treatment groups. Each row of the heatmap represents the z-score transformed log 2 values of one differentially expressed gene across all samples (blue, low expression; red, high expression). (B) Volcano plots of significantly differentially expressed genes, (-log10 ≤ 0.05 and log2 ≥ 1.5; red, up-regulated; blue, down-regulated). The top 30 upregulated or down-regulated genes are denoted.

**Table 1.**

| **Entrez gene** | **Gene Symble** | **Fold-change** | **Regulation** |
| --- | --- | --- | --- |
| Top 10 down-regulated DEGs |  |  |  |
| 27019 | DNAI1 | -2.915444524 | Down |
| 11045 | UPK1A | -2.905284028 | Down |
| 79977 | GRHL2 | -2.703445298 | Down |
| 102724219 | LOC102724219 | -2.608568444 | Down |
| 2994 | GYPB | -2.444313829 | Down |
| 284525 | SLC9C2 | -2.320280303 | Down |
| 347168 | OR1J1 | -2.283834562 | Down |
| 6540 | SLC6A13 | -2.211703973 | Down |
| 290 | ANPEP | -2.196158711 | Down |
| 434 | ASIP | -2.185374364 | Down |
| Top 10 up-regulated DEGs |  |  |  |
| 100169851 | PATE3 | 2.97403579 | up |
| 100129520 | TEX13C | 2.29248125 | up |
| 538654 | TMEM110 | 2.241526861 | up |
| 440087 | SMCO3 | 2.201838731 | up |
| 150356 | CHADL | 2.03630614 | up |
| 105371191 | LOC105371191 | 1.979453368 | up |
| 112267874 | LOC112267874 | 1.923565413 | up |
| 346 | APOC4 | 1.894025701 | up |
| 3681 | ITGAD | 1.871475957 | up |
| 79585 | CORO7 | 1.842316162 | up |

### Enrichment analysis of DEGs between the control and SI2-treatment MM cells

To analyze the functional pathways and mechanism during the SI2 treatment of MM, we performed KEGG pathway and GO categories analysis (Figure 2). Our study showed the top three KEGG pathways with P value < 0.05, including “Neuroactive ligand-receptor interaction”, “Hemaotopoietic cell lineage”, and “Complement and coagulation cascades”.

**Figure 2.**
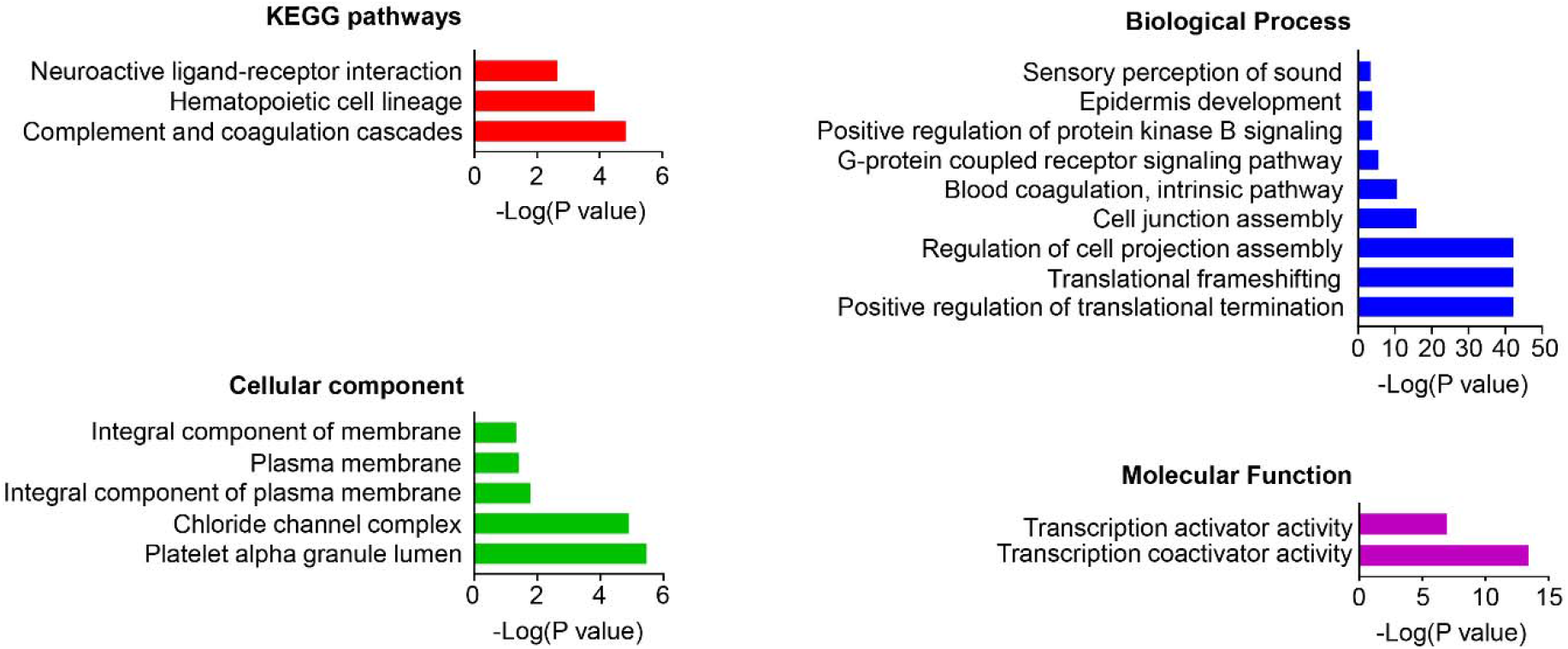
KEGG and GO analyses. The chart of enrichment results grouped by clusters (labeled on the left-hand side of each panel). The x axis corresponds to fold enrichment values, while the y axis indicates the enriched pathways. The bar length indicates the number of differentially expressed genes (DEGs) in the given pathway.

We further analyzed the GO enrichments by three categories. We identified nine significant items in biological processes including “Sensory perception of sound”, “Epidermis development”, “Positive regulation of protein kinase B signaling”, “G-protein coupled receptor signaling pathway”, “Blood coagulation, intrinsic pathway”, “Cell junction assembly”, “Regulation of cell projection assembly”, “Translational frameshifting”, and “Positive regulation of translational termination”. We also identified five cellular components including “Integral component of membrane”, “Plasma membrane”, “Integral component of plasma membrane”, “Chloride channel complex”, and “Platelet alpha granule lumen”. We further identified two main molecular functions in GO including “Transcription activator activity” and “Transcription coactivator activity”.

### PPI network and Module analysis

We created the PPI network to further study the relationships of DEGs. We set the criterion of combined score (> 0.7) and constructed the PPI network by using the 345 nodes and 188 interactions between the control and SI2-treatment MM cells. The top two significant modules were selected to depict the functional annotation of genes (Figure 3).

**Figure 3.**
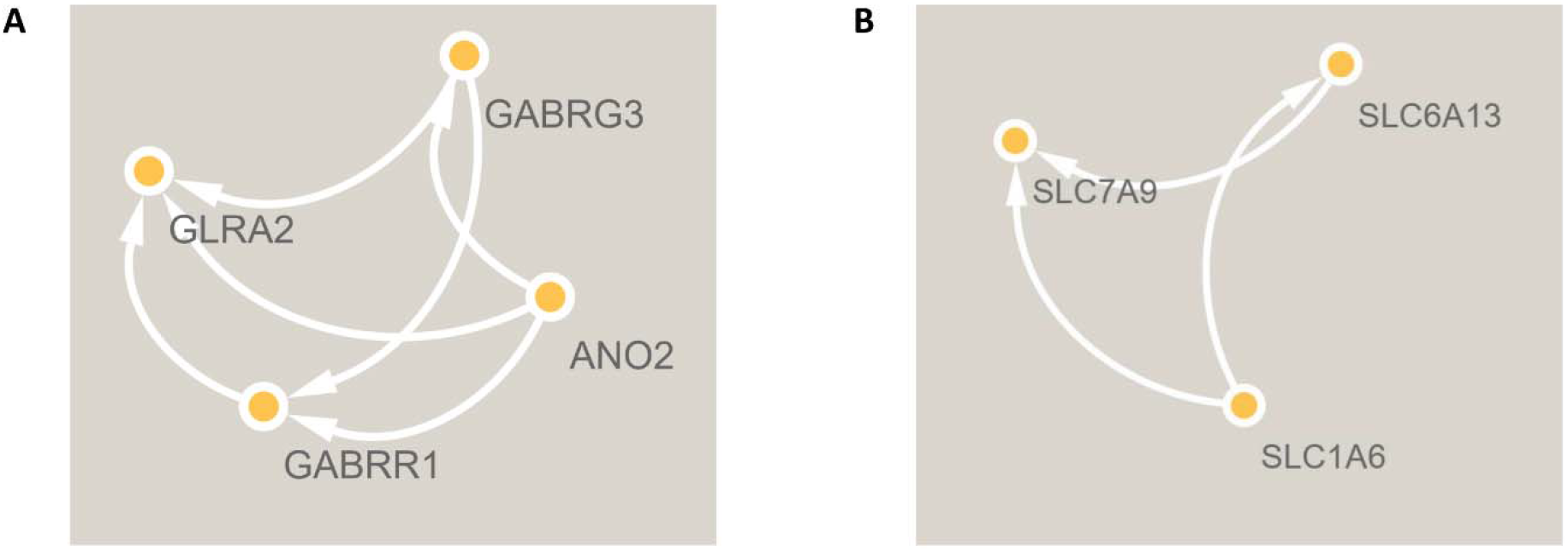
The protein-protein interaction (PPI) network analysis of differentially expressed genes using STRING database. The 384 differentially expressed genes were input into STRING database for PPI network analysis. The main cluster 1 (A) and cluster 2 (B) were analyzed by MCODE in Cytoscape, and the nodes in each cluster were input into STRING database to obtain the PPI subnetworks.

To further study the signaling pathways, we also performed the Reactome analysis (Figure 4 and Table S1). The top signaling pathways were selected (P < 0.1) including “Formation of Fibrin Clot”, “Intrinsic Pathway of Fibrin Clot Formation”, “GP1b-IX-V activation signalling”, “Interaction with Cumulus Cells and The Zona Pellucida”, “Defective factor IX causes hemophilia B”, “Physiological factors”, “RUNX1 regulates genes involved in megakaryocyte differentiation and platelet function”, and “IL-6-type cytokine receptor ligand interactions”.

**Figure 4.**
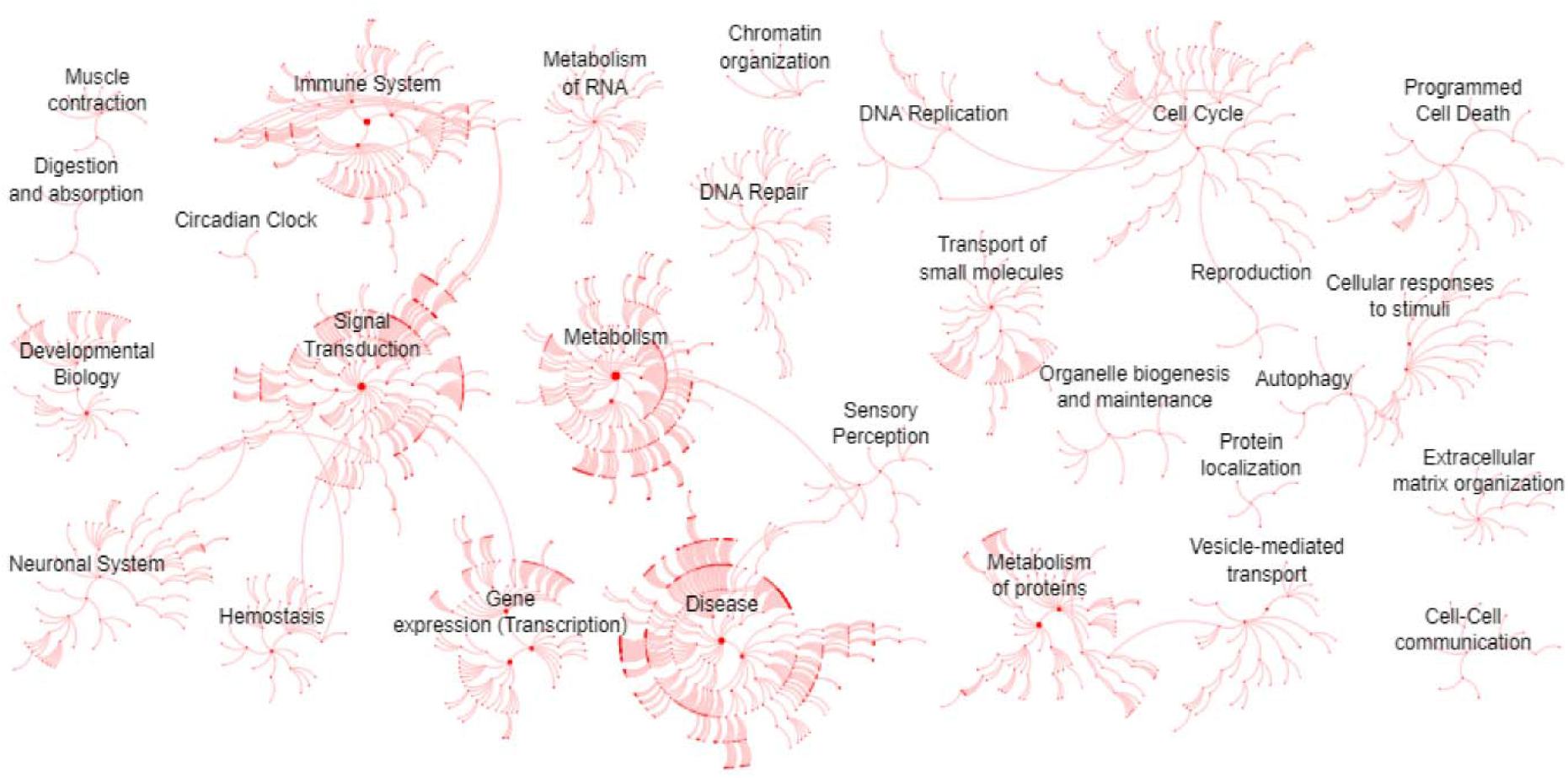
Reactome map indication of the distribution of biological processes in which significant differentially expressed genes identified via mRNA sequencing listed in Figure 1 are involved.

## Discussion

Small-molecule inhibitors for blocking PPI to eliminate activities of target proteins have shed light on methods to exploit strategies for cancer therapy^16–18^. SI2 was found to significantly repress cancer growth through a direct binding with SRC-3, which can reduce the protein expression and transcriptional activities of SRC-3 without affecting the cellular viability^19^. Moreover, SI2 also showed no obvious toxicity and inhibition of lung tumor by repressing the number of cancer-like cells^20^. Thus, this study is mainly focused on the gene function and mechanism of SI2 inhibiting the MM and to find the key pathway for SI2 to inhibit cancer.

MM was reported to affect blood composition and a number of MM patients have been found Bence-Jones proteinemia during complement-fixation tests^21^. In our study, the KEGG results showed the complement and coagulation cascades and Hematopoietic cell lineage were also affected during the treatment. G-protein-coupled receptor signaling pathways and regulators of G protein signaling (RGS) signaling pathways play critical roles in physiological and pathological processes such as bone formation, arthritis, cancer and pain^22–24^. Kodandaram Pillarisetti et al showed that the G-protein-coupled receptor class 5 member D (GPRC5D) is expressed in MM and can eliminate the MM cells^25^. We also found the use of SI2 treatment could affect the G-protein coupled receptor signaling pathway in biological processes. A number of reports found that several transvectors were changed in MM. NF-κB is the critical factor during the inflammatory diseases^26, 27^, which is found to regulate the MM autonomy from the bone marrow microenvironment^28^. In molecular function analysis, we found the transcription activator activity and transcription coactivator activity were both affected by the treatment of SI2, which suggested that targeting the transcription factor such as NF-kB, p53, clock may inhibit the tumorigenesis.

In this study, we identified the PPI network and potential genes may be the key factors during the treatment of MM or other cancers. Phosphatase of regenerating liver-3 (PTP4A3/PRL-3) was found to activate the Src and Src family kinase (SFK) members such as LYN and HCK in MM cells^29^. Circadian gene clocks control most cell functions such as metabolism, proliferation, and differentiation^30–36^. Genome-wide interaction analysis in multiple myeloma showed the circadian rhythm plays a key role in regulating the development of bone marrow microenvironment^37^. Interestingly, the serum erythropoietin showed the circadian rhythm exists in the MM patients, which suggests the multiple circadian rhythm system may be involved in the regulation of MM metabolism^38^. Erin Stashi found the SRC-2 is a potential coregulator of BMAL1:CLOCK, which targets itself with BMAL1:CLOCK in a feedback loop^39^. Jiekai Yu et al found the Kininogen 1 is a serum protein marker for colorectal cancer^40^. Jinfang Xu et al found the forced expression of Kininogen-1 represses the glioma cells proliferation and promotes apoptosis^41^. Proteomic data showed the Kininogen 1 is closely related to the development of MM^42^. GPCR and RGS signaling pathways play key roles in maintaining the cell functions and metabolisms^43–46^. Class I phosphoinositide 3-kinase (PI3K), also known as PI3Kgamma, links GPCR signaling pathways to regulate cellular movement, adhesion, and secretion^47^. Xiaofeng Wang et al found the exosomal miR-301a controls M2 Macrophage Polarization via PTEN/PI3Kγ to drive the pancreatic cancer metastasis^48^. Importantly, PI3Kγ plays a critical role in mediating the pro-tumoral signaling in MM microenvironment^49^. Herdman C et al found that deficiency of ESYT3 does not impact on the mouse development in vivo but affects the cell migration in vitro^50^. The endoplasmic reticulum (ER) plays a protective role in maintaining proteostasis due to its involvement in protein quality control^51–53^. ESYT3 was found to affect hypothalamic ER stress to regulate the cell functions^54^. Histamine N-methyltransferase (HNMT) is the enzyme of histamine, which has a key function in regulating autoimmune diseases^55^. KCNH2 is important for prevention and treatment of potassium channels related diseases and cancers^56, 57^. K D Sørensen et al found 5-aza-2’-deoxycytidine promoted ANPEP expression in prostate cell lines, which suggested there is an epigenetic silencing^58^. Y-F Li et al found the expression of MYOZ2 is significantly higher than that in cancer-adjacent tissues, suggesting it has a potent relationship with cancer^59^. GABRG3 was found to be relevant with the inflammatory pain and nervous system diseases^60^. Alexander Cox et al found the Otoferlin is a prognostic marker in renal carcinoma patients^61^.

In conclusion, we provided the knowledge for the treatment of MM with SI2. The complement and coagulation cascades and the transcription activator activity are the main biological processes during the treatment of SI2 in MM cells. Our future studies will focus on exploring the upstream and downstream of these significant pathways. Our study may facilitate the drug development of MM.

**Table 2.** Top ten genes demonstrated by connectivity degree in the PPI network.

| Gene Symbol | Gene title | Degree |
| --- | --- | --- |
| SRC | SRC proto-oncogene, non-receptor tyrosine kinase | 19 |
| KNG1 | Kininogen 1 | 9 |
| PI3KCG | Phosphatidylinositol-4,5-bisphosphate 3-kinase catalytic subunit gamma | 7 |
| ESYT3 | Extended synaptotagmin 3 | 5 |
| HNMT | Histamine N-methyltransferase | 5 |
| KCNH2 | Potassium voltage-gated channel subfamily H member 2 | 5 |
| ANPEP | Alanyl aminopeptidase, membrane | 4 |
| MYOZ2 | Myozenin 2 | 4 |
| GABRG3 | Gamma-aminobutyric acid type A receptor subunit gamma3 | 4 |
| OTOF | Otoferlin | 4 |

## Author Contributions

Hanming Gu: Conceptualization, Methodology, Writing-Reviewing and Editing.

